# Agonistic anti-CD40 monoclonal antibody improves the antitumor efficacy of a clinical chemo-immunotherapy regimen in a refractory mouse neuroblastoma model

**DOI:** 10.64898/2026.11.25.769196

**Authors:** Alexander L. Rakhmilevich, Joel Matthews, Mildred AR. Felder, Alexa K. Heaton, Anqi Gao, Adithyan Unni, Katherine C Delcore, Benjamin B. Horwitz, Colleen Carlisle, Cole Witt, Nikolai P. Hess, Jen Zaborek, Melissa Skala, Huy Dinh, Amy K. Erbe, Paul M Sondel

**Affiliations:** Department of Pediatrics, University of Wisconsin, Madison, WI; Department of Radiation Medicine, University of Wisconsin, Madison WI; Morgridge Institute for Research, Madison WI; Department of Biomedical Engineering, University of Wisconsin, Madison WI; Department of Oncology, University of Wisconsin, Madison WI; Department of Biostatistics and Medical Informatics, University of Wisconsin, Madison WI

**Keywords:** Neuroblastoma, chemotherapy, immunotherapy, anti-CD40 antibody, preclinical

## Abstract

**Background:** The current standard regimen for relapsed-refractory high-risk neuroblastoma (HR-NB) is the combination of chemotherapy agents -**<u>T</u>**emozolomide and **I**rinotecan - and anti-**<u>G</u>**D2 antibody dinutuximab (TIG). This treatment results in an overall response for ∼40% of patients, but the remaining 60% are resistant to TIG. We hypothesized that agonistic anti-CD40 monoclonal antibody (aCD40), which activates antitumor NK cells and macrophages as we found in previous studies, will improve the efficacy of TIG.

**Aim:** To test the combination of aCD40 and TIG in a mouse HR-NB model that is resistant to TIG.

**Methods:** For these studies, we used a clinically relevant, immunologically cold, HR-NB murine model, 9464D-GD2. Syngeneic mice were intradermally injected with the tumor cells. When tumors were established, mice were randomized and treated with TIG and/or aCD40. Tumor growth and mouse survival were followed. Mice that rejected their tumors were rechallenged to test immunological memory. Changes in immune cells were evaluated by flow cytometry and single-cell RNA sequencing (scRNAseq).

**Results:** Similar to ∼60% of patients with relapsed/refractory HR-NB, mice bearing 9464D-GD2 tumors did not respond to TIG. Adding aCD40 to TIG induced complete tumor regression in most mice bearing small tumors and extended survival, with many mice remaining tumor-free. The combination of aCD40 and TIG was more effective than aCD40 alone. The antitumor effect of TIG+aCD40 was not inhibited in T cell- or neutrophil-depleted mice but was substantially reduced in NK cell- and macrophage-depleted mice, indicating a role for both NK cells and macrophages in the antitumor effect. Involvement of these cells was consistent with flow cytometry and scRNAseq analyses. Mice treated with TIG+aCD40 that remained tumor-free were rechallenged with 9464D-GD2 cells and showed significantly reduced tumor growth compared to naïve mice, indicating a memory response. This memory response was reduced in T cell-depleted mice, indicating the involvement of T cells in memory.

**Conclusion:** These results show that while TIG alone was ineffective, a combination of aCD40 and TIG induced substantial tumor eradication in the 9464D-GD2 tumor model, suggesting that aCD40 may warrant future testing as an addition to the current standard TIG regimen for children with relapsed-refractory HR-NB.

**What is already known on this topic:**

- For children with relapsed or refractory neuroblastoma, the addition of anti-GD2 mAb to chemotherapy consisting of temozolomide and irinotecan is much more effective than chemotherapy alone. Despite this progress, about 60% of patients are resistant to this combined therapy.

**What this study adds:**

- We show that adding agonistic CD40 antibody activates NK cells and macrophages and dramatically improves the efficacy of the temozolomide/irinotecan/anti-GD2 regimen in this mouse model of MYCN-driven, refractory neuroblastoma.

**How this study might affect research, practice or policy:**

- Our findings suggest that the addition of anti-CD40 antibody may warrant future clinical testing to determine if it improves the efficacy of a current standard regimen for relapsed high-risk neuroblastoma in children.

## Introduction

Neuroblastoma (NB) is the most common extra-cranial solid tumor in children. About 50% of newly diagnosed patients with NB have clinical and tumor-related features indicative of high-risk (HR) disease, associated with poor prognosis. Adding anti-GD2 monoclonal antibody (mAb) immunotherapy to the post-consolidation maintenance improves outcomes for these patients.^1^ However, ∼50% of the patients are either refractory to the initial treatment or relapse following it.^2–4^ For patients with relapsed or refractory NB, adding anti-GD2 mAb to temozolomide (TEM) and irinotecan (IRI) chemotherapy was much superior to chemotherapy alone.^5,6^ Despite this progress, ∼60% of relapsed/refractory patients were resistant to this chemo-immunotherapy.^5–7^ Besides early studies using anti-GD2 CAR-T cells, there are currently no effective treatment options for patients who have failed salvage therapy with chemotherapy and anti-GD2 mAb.^8,9^ A more effective regimen for patients with relapsed-refractory HR-NB is needed.

One activator of innate and adaptive immunity is agonistic anti-CD40 antibody (aCD40), which can induce antitumor effects in mice and in some cancer patients.^10,11^ Multiple aCD40 agonistic antibodies have been used in clinical trials, alone or combined with other treatments, showing antitumor activity in some, but not all trials.^12–15^ A Phase-I trial has been completed in children with CNS tumors. ^12^ One early trial demonstrated regression in 4 of 21 patients with pancreatic cancer.^11^ This clinical activity of aCD40 is consistent with our earlier demonstration of antitumor effects with aCD40 via NK cell- and macrophage activation in mice.^16–18^ We have recently found that adding aCD40 to anti-GD2 mAb delays tumor growth in mice bearing 9464D-GD2 tumors; during that response immune cells, including macrophages and NK cells, are increased within the TME.^19^ Yet, tumors eventually grow out, indicating additional therapeutic measures are needed. We separately showed that a combination of chemotherapy (cyclophosphamide, doxorubicin and vincristine) and immunotherapy including aCD40 converted M2 pro-tumor macrophages into M1 antitumor-effector macrophages, and demonstrated enhanced antitumor efficacy in mouse models including 9464D NB,^20^ suggesting that aCD40 could be combined with <u>T</u>EM/IRI/anti-<u>G</u>D2 (TIG).

In this study, we show that the immunologically cold 9464D-GD2 NB with a low mutation burden^21^ is resistant to TIG; however, addition of CD40 yielded tumor eradication in mice bearing small tumors and substantial tumor growth inhibition in mice with larger tumors.

## Materials and Methods

### Mice

Male and female 7–12 week-old C57BL/6 mice obtained from Taconic Farms (Germantown, NY) or bred at the University of Wisconsin-Madison (UW) animal facility were used according to the *Guide for Care and Use of Laboratory Animals* (NIH publication 86-23, 1985). The macrophage mCherry reporter mouse model was made using Cre-lox methods by crossing Lyz2-Cre mice (Jax 004781) with floxed diphtheria toxin receptor (DTR)-mCherry mice under the control of the *Csf1r* promoter (Jax 024046) at UW as previously described.^22^ Lyz2-Cre homozygous / Csf1r-DTRmCherry hemizygous mice were bred, genotyped, and used for these experiments.

### Tumor cell line

Mouse 9464D-GD2 Cl.7.2 neuroblastoma, syngeneic to C57BL/6 mice, having a low tumor mutation burden (1.6 mutations/megabase) and expressing minimal MHC-I even after stimulation with IFN-γ, was grown and mycoplasma-tested as described.^21,23^

### Reagents

Agonistic aCD40 was obtained from the FGK 45.5 hybridoma (F. Melchers, Basel, Switzerland) and prepared as described.^16,17^ TEM was obtained from Sigma-Aldrich, St. Louis, MO and Medchemexpress LLC, Monmouth Junction, NJ. IRI was obtained from Apotex, Weston, FL. The murine anti-GD2 mAb (14G2a) was obtained from Brunswick BioTechnetics through the NCI BRB Repository and BioXCell, Lebanon, NH. Diphtheria toxin (DT) was obtained from Sigma-Aldrich.

### *In vivo* tumor models

Intradermal (i.d.) tumors were established by injecting 2×10^6^ 9464D-GD2 cells in 0.1 ml of PBS into the flank or abdomen.^21^ Tumor diameters were measured with digital calipers, and tumor volume (mm^3^) was calculated as ½ x tumor length x tumor width^2^. On various days prior to treatment start (based on the desired size of tumors at the start of treatment), tumor-bearing mice were randomized to achieve groups with similar tumor volumes. Treatments and tumor measurements were performed in blind fashion. The first day of treatment is defined as “day 0”. TEM (64 *μ*g/dose) and IRI (32 *μ*g/dose) were injected intraperitoneally (i.p.) daily on days 0-4. Anti-GD2 antibody (100 *μ*g/dose) was injected i.p. on days 0 and 4. aCD40 (500 *μ*g/dose) was injected i.p. on day 1. For T cell and NK cell depletion, anti-CD4, clone GK1.5 (200 *μ*g/dose), anti-CD8, clone 2.43 (100 *μ*g/dose), anti-NK1.1., clone PK136 (100 *μ*g/dose), or rat IgG (300 *μ*g/dose) was injected i.p. on days -1, 3, 6, and 10. To deplete neutrophils, anti-Ly6G mAb, clone 1A8 (500 *μ*g/dose) or control rat IgG (500 *μ*g/dose) was injected i.p. on days 0, 1, 3, 5, and 7. Depleting antibodies were purchased from BioXCell, and rat IgG was purchased from Sigma-Aldrich. For macrophage depletion, 9464D-GD2 tumor cells were injected i.d. into DTR-mCherry-*Csf1r* mice ∼26 days before treatment. Each randomized group included male and female mice. DT (100 ng in 200 μL PBS) was administered i.p. once daily on days -1, 1, and 3. For rechallenge experiments, mice that rejected 9464D-GD2 tumors following TIG + aCD40 and remained tumor-free were rechallenged three months later i.d. with 2×10^6^ 9464D-GD2 tumor cells in the opposite flank. Naïve control mice were injected with the same tumor cells.

### Flow cytometry

C57BL/6 mice bearing 9464D-GD2 NB were treated with TIG +/- aCD40. On day 5 unless stated otherwise, tumors were harvested and disaggregated into single-cell suspension and stained to identify the different cell populations and subpopulations, as described previously,^23,24^ and detailed in **Supplementary Materials and Methods**.

### scRNA-seq sample collection, library preparation, and sequencing

Treatments of 9464D-GD2 tumor-bearing mice were performed as described above. Six days after the treatment started, seven to ten tumors for each treatment group were resected and pooled. Tumors were dissociated, isolated, processed, and analyzed for scRNAseq, as previously reported,^25^ and as detailed in the **Supplementary Materials and Methods**. Single-cell RNA-seq data have been deposited at GEO and are publicly available (GEO code: GSE310495). Any additional information required to reanalyze the data reported in this paper is available from the lead contact upon request.

### Statistical analysis

Average group tumor volumes are plotted showing mean +/- SEM. Tumor volume plots were summarized by time-weighted average (area under the volume-time curve, calculated using the trapezoidal method). Time-weighted averages were compared between treatment groups overall by Kruskal-Wallis tests. In tumor growth plots, ratios shown are tumor-free mice/total mice in that group. If significant by Kruskal-Wallis test or if only two groups were determined to be compared a priori, pairwise comparisons were conducted using Mann-Whitney tests. Survival data were plotted using Kaplan-Meier methods and analyzed using log-rank comparisons. Flow cytometry results are plotted as mean +/- SD, with each symbol representing results from one mouse/sample. Flow cytometry data were analyzed by the Mann-Whitney-Wilcoxon test. Complete response rates were analyzed overall with chi-square tests (when comparing three or more groups). If significant by chi-square tests or if only two groups were determined to be compared a priori, pairwise comparisons were conducted using two-sample tests of proportions. P values < 0.05 were considered significant. All experiments were repeated twice unless stated otherwise. For all graphs \**p* < 0.05, \*\**p* < 0.01, \*\*\**p* < 0.001, \*\*\*\**p* < 0.0001.

## Results

### 9464D-GD2 HR-NB resistance to TIG is overcome by addition of aCD40

We previously used 9464D-GD2, a MYCN-driven, immunologically cold, low mutation burden mouse HR-NB that simulates human HR-NB^5–7,21^. To determine its sensitivity to TIG, mice bearing 9464D-GD2 tumors with different tumor volumes (4, 8, or 16 mm^3^) were treated with a 5-day course of TIG (**Figure 1A**). The results show that even barely palpable tumors (4 mm^3^) were resistant to TIG; no tumor regression was observed in any TIG-treated mice (**Figure1B**). Based on our previous data showing that a combination of immunotherapy including anti-CD40 and chemotherapy (cyclophosphamide, doxorubicin and vincristine) had a greater antitumor effect against 9464D NB than immunotherapy or chemotherapy alone^20^, we hypothesized that adding aCD40 to TIG would improve antitumor efficacy. Indeed, our results show that whereas TIG was ineffective against ∼17 mm^3^ 9464D-GD2 tumors, a combination of TIG and aCD40 induced complete tumor regression (**Figure 1C,D**) and tumor-free survival (**Figure 1E**) in most of the treated mice. The mean tumor growth curves in **Figure 1D** also show that TIG alone, without the addition of aCD40, does not even slow the growth of these small 17 mm^3^ 9464D-GD2 tumors compared to the no treatment (NT) control.

**Figure 1.**
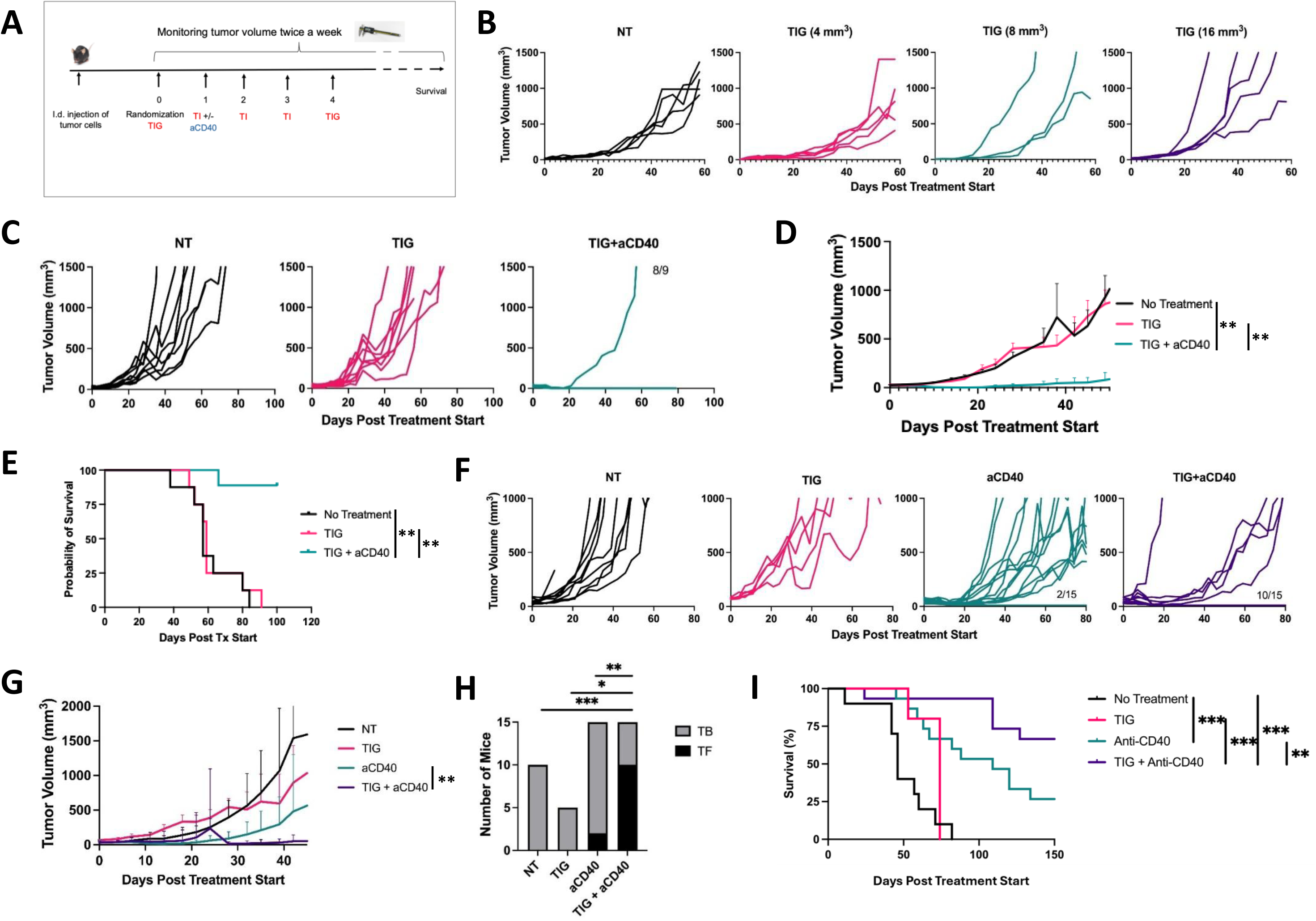
9464D-GD2 NB is resistant to TIG but sensitive to TIG+aCD40. **A.** Treatment schema. **B**. C57BL/6 mice were injected i.d. with 9464D-GD2 tumor cells in the right flank and were treated with TIG starting when mean tumor volume reached 4, 8, or 16 mm^3^. NT = No treatment. Each line represents a tumor volume from a separate mouse. **C-E**. Mice bearing 9464D-GD2 tumors received TIG+/-aCD40 when tumors were 17 mm^3^. Tumor growth in individual mice **(C)** and means +/-SEM **(D**) are shown. Survival (**E**) was followed. At 100 days post-treatment start, 8 out of 9 mice in the TIG+ aCD40 group were alive and tumor-free. **F-I**. C57BL/6 mice bearing 9464D-GD2 tumors received TIG, aCD40, TIG+aCD40, or NT when the tumors averaged 17 mm^3^. **F**. Growth of individual tumors. **G.** Mean +/- SEM for the 4 groups shown in **F. H.** Numbers of tumor-bearing (TB) mice (including dead mice due to progressive tumor) and tumor-free (TF) mice by the end of the experiment. **I**. Survival of mice. Combined data from three independent experiments are shown. \**p* < 0.05, \*\**p* < 0.01, \*\*\**p* < 0.001, \*\*\*\**p* < 0.0001. Numbers in **C** indicate the ratio of tumor-free mice to total mice per group.

Next, we asked if the antitumor effects induced by the combination of TIG and aCD40 are better than the effects of aCD40 alone. The results show that the combination of TIG and aCD40 induced statistically greater effects than either TIG or aCD40 alone, as seen in suppression of tumor growth (**Figure 1F,G**), proportion of tumor-free mice (**Figure 1H**), and survival (**Figure 1I**). Treatment with aCD40 or TIG+aCD40 resulted in transient toxicity as shown by reduced mouse weight and elevated liver enzymes, which peaked around day 3 (2 days after aCD40 treatment), remained significantly elevated compared to untreated mice on day 5, and reached control levels by day 9 (**Supplementary Figure 1**).

To understand why aCD40 + TIG was more effective than aCD40 alone, we asked if TIG might induce upregulation of CD40 molecules on macrophages, as we previously showed an upregulation of CD40 on macrophages by prior treatment with CpG.^16–21^ Indeed, peritoneal macrophages expressed increased CD40 on their surface 1 day after i.p. TIG treatment, although this increase was not observed on the surface of intratumoral macrophages (**Supplementary Figure 2**).

### Role of immune cells in the antitumor effect of TIG+aCD40

#### Role of T and NK cells

To determine if an immune system is necessary for the antitumor effect of TIG+aCD40, we compared TIG+aCD40 efficacy simultaneously in immunocompetent and immunodeficient NRG mice (lacking T, B, and NK cells, with reduced macrophage function). We observed antitumor efficacy as shown by tumor regression and increased number of tumor-free mice in immunocompetent C57BL/6 mice treated with aCD40 and TIG+aCD40 (**Figure 2A,B**), but no tumor regression was observed with the same therapies in NRG mice (**Figure 2C,D**), suggesting that immune cells are activated by TIG+aCD40 and are essential in eradicating 9464D-GD2 tumors with this treatment.

**Figure 2.**
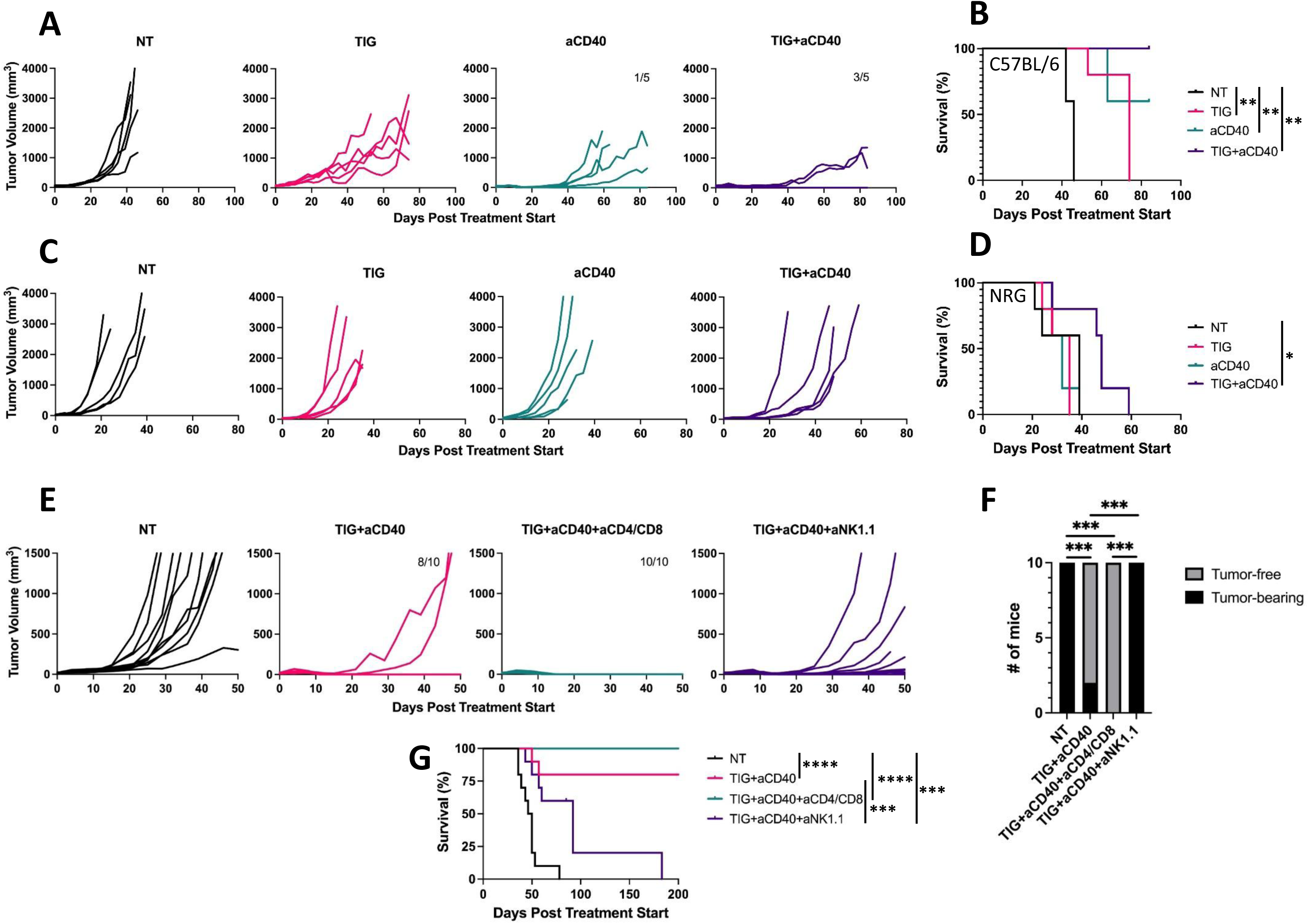
Role of T cells and NK cells in the TIG+aCD40-induced antitumor effect. C57BL/6 mice (**A**,**B**) and NRG mice (**C**,**D**) bearing 9464D-GD2 tumors were treated with TIG, aCD40, TIG+aCD40, or NT. **A,C**. Growth of individual tumors. **B,D**. Survival of mice. One of two independent experiments for each strain is shown. **E-G**. Immune cell depletion studies in C57BL/6 mice. **E**: individual mouse tumor curves; **F**: Numbers of TB and TF mice by the end of the experiment; and **G**: survival following treatment with TIG+aCD40 and either rat IgG, anti-CD4 + anti-CD8, or anti-NK1.1 antibodies. Combined data from two independent experiments are shown.

To directly address the role of T cells and NK cells in the antitumor effect of TIG+aCD40, we performed antibody-depletion experiments. While depletion of CD4^+^ and CD8^+^ T cells did not interfere with the antitumor effect of TIG+aCD40 against 9464D-GD2 NB (nearly all mice in both groups became and remained tumor-free), depletion of NK cells significantly suppressed the antitumor effect of TIG+aCD40; none of the NK-depleted mice remained tumor-free and the survival was significantly reduced compared to TIG+aCD40-treated mice (**Figure 2E,F,G**). However, the initial tumor growth suppression (**Figure 2E**) and prolongation of survival compared to NT mice (**Figure 2G**) were evident even in NK cell-depleted mice, suggesting that an additional mechanism contributed to the antitumor effect in the absence of NK cells.

*Role of neutrophils and macrophages*. The statistically significant prolongation of survival in NRG mice following TIG+aCD40 treatment (**Figure 2D**), and in C57BL/6 mice depleted of NK cells (**Figure 2G**), suggest that some other immune/inflammatory cells that remain in these mice may be activated by the combined treatment to suppress 9464D-GD2 growth. We thus evaluated the role of neutrophils and macrophages. Neutrophils can be antitumor effector cells following aCD40 administration for some tumors,^25–27^ but have also been shown to have a negative effect on immunotherapy outcome in other settings,^28^ and in 9464D-GD2-bearing mice treated with anti-GD2 + anti-CD40 + epigenetic modifier inhibitors.^19^ Thus we tested if neutrophil depletion with anti-Ly6G mAb would inhibit the antitumor efficacy of TIG+aCD40. The results show that neutrophil depletion did not reduce the antitumor effect of TIG+aCD40 but rather significantly augmented it (**Figure 3A-D**), suggesting that neutrophils may exert immunosuppressive rather than antitumor function in C57BL/6 mice bearing 9464D-GD2 treated with TIG+aCD40.

**Figure 3.**
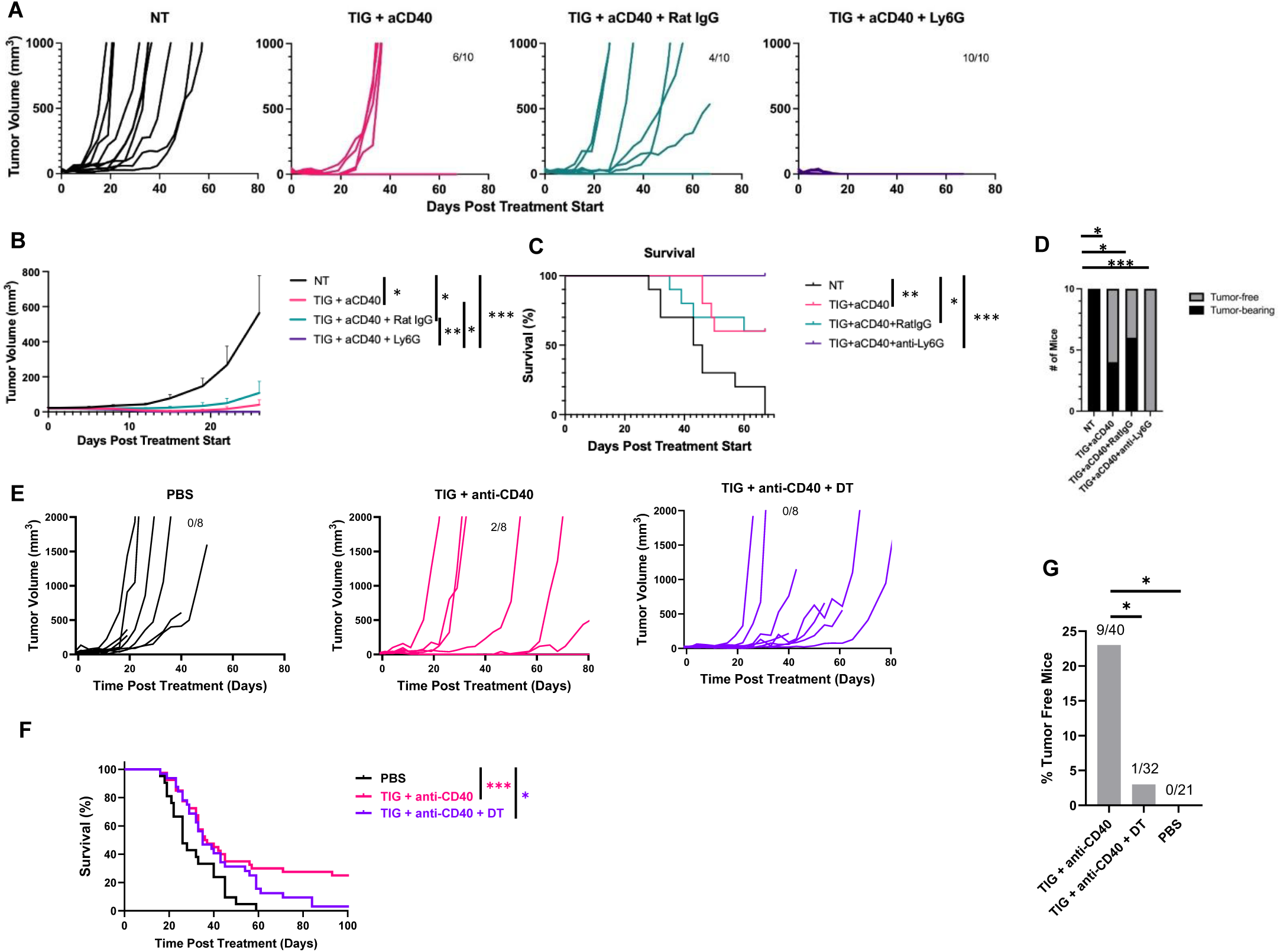
Role of neutrophils and macrophages in the TIG+aCD40-induced antitumor effect. C57BL/6 mice (**A-D**) and Csf1r-DTR/mCherry mice (**E-H**) bearing 9464D-GD2 tumors were treated with TIG+aCD40, PBS or NT. **A-D.** Mice were depleted of neutrophils with anti-Ly6G mAb or rat IgG (control). Individual tumor growth curves (**A**), means +/-SEM **(B**), survival (**C**) and number of tumor-free mice (**D**) are shown. Data from two combined experiments are shown. **E-G**. Csf1r-DTR/mCherry mice were transiently depleted of macrophages with DT on days -1, 1, and 3. TIG + aCD40 treatment was given as shown in Figure 1A. Individual tumor growth curves (**E**) of one representative experiment of 4 are shown (*n*=8). Survival (**F**) and number of tumor-free mice by the end of the experiments on day 100 (**G**) are shown as the combined data of four independent experiments (n=21-40 total per group).

Based on our previous findings that aCD40 activates antitumor macrophages^16–19^ and has anti-tumor activity in vivo in mice bearing tumors of the parent 9464D cell line^20^, and based on the reduction of tumor growth in TIG+aCD40–treated NRG mice and NK-depleted C57BL/6 mice (**Figures 2D and 2G**), we hypothesized that TIG+aCD40 can activate macrophages to kill 9464D-GD2 tumor. To deplete macrophages *in vivo*, we used DT treatment in transgenic Csf1r-DTR/mCherry macrophage reporter mice. Flow cytometry confirmed that following injection of DT, macrophages were markedly depleted (average 95% depletion, n=3) in the peritoneal cavity, and incompletely depleted (average 57% depletion, n=3) in 9464D-GD2 tumors (**Supplementary Figure 3**). When macrophages were depleted during TIG+aCD40 treatment, the antitumor efficacy was significantly reduced (**Figure 3E-G**), suggesting an important role of macrophages in driving this antitumor response.

#### Flow cytometric analysis of immune cells in the tumor microenvironment (TME)

Flow cytometric analysis of immune cells in the TME showed that TIG+aCD40 treatment resulted in reduced CD4^+^ T cells, increased NK cells and neutrophils (Ly6G^+^CD11b^+^), and no significant change in CD8^+^ T cells, Tregs (CD4^+^CD25^+^FoxP3^+^), macrophages (F4/80^+^CD11b^+^) and myeloid-derived suppressor cells (CD11b^+^Gr-1^+^) (**Figure 4A,B**). Additionally, we observed a significant decrease in the percentage of CD206^+^ macrophages (CD206 is a mannose receptor expressed on M2 macrophages) while the percentage of MHC class II macrophages (M1 marker) did not change (**Figure 4A**).

**Figure 4.**
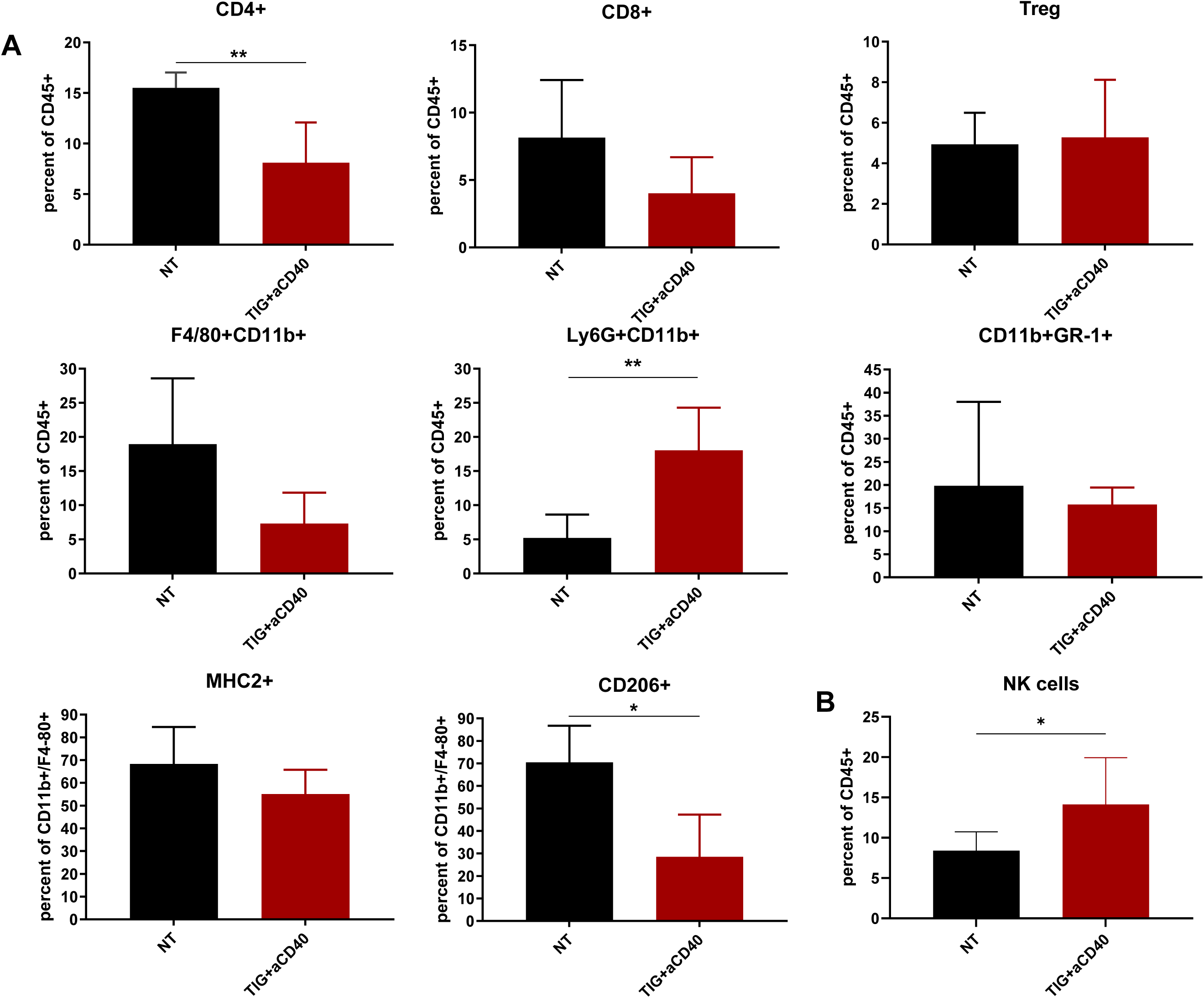
Effect of TIG+aCD40 treatment on the TME. C57BL/6 mice bearing 9464D-GD2 tumors were treated with TIG+aCD40 or NT as in Figure 1A. On day 5, tumors were harvested and analyzed by flow cytometry. Cells were gated on live single CD45^+^ cells. **A**. Mean +/-SD percentages are shown for T cells (CD4^+^, CD8^+^ T cells and Tregs), macrophages (CD11b^+^F4/80^+^), neutrophils (CD11b^+^Ly6G^+^), and myeloid-derived suppressor cells (CD11b^+^Gr1^+^). In the last two graphs, the cells were gated on macrophages (CD11b^+^ F4/80^+^), and the percentages of MHC class II (MHC2^+^) and CD206^+^ cells are shown. Data from one of two similar experiments are shown (n=5). **B**. Mean +/-SD percentages are shown for NK cells using the combined data from two similar experiments (n=8).

#### Single-cell transcriptomic analyses of immune cells in the 9464D-GD2 TME

TIG caused no or little benefit in survival or tumor growth inhibition compared to mice receiving no treatment, while adding aCD40 to TIG caused substantial tumor growth inhibition, improved survival, and (depending on tumor size at treatment initiation) a substantial fraction of mice that became and remained tumor-free (**Fig. 1B-D**). Thus, we wanted to analyze the gene expression of the cells potentially involved in the immunotherapeutic benefit that we found in tumors from mice treated with TIG + aCD40 vs. tumors from mice treated with TIG alone. Furthermore, in this 9464D-GD2 model, aCD40 treatment alone caused a benefit in survival and tumor growth inhibition compared to no treatment, but adding TIG to aCD40 caused further benefit (**Fig. 1F-I**). Thus, we also analyzed the gene expression of the immune cells, comparing results from mice treated with aCD40 alone vs. mice treated with TIG+aCD40. As our depletion analyses have shown a prominent role for both NK cells (**Figure 2E-G**) and macrophages (**Figure 3E-G**) in the benefit found by adding aCD40 to TIG, while T cells were not required for the response to TIG+aCD40 (**Fig. 2E-G**), we have focused our scRNAseq analyses on NK cells and macrophages.

To evaluate tumor-infiltrating immune cell changes, we performed scRNA-seq of CD45^+^ cells from the TME of 9464D-GD2-bearing mice on day 6 following treatment with TIG, aCD40, or TIG+aCD40. scRNA-Seq captured 15,020 immune cells across all major immune cell type compartments in all three treatment regimens (**Figure 5A, Supplementary Figure 4**). While diversity of T cell subsets was observed for all 3 regimens, we did not observe significant differences in frequencies between the 3 regimens (**Supplementary Figure 4**), with a notable ∼30% of regulatory T cells remaining similar across all treatments. Within the myeloid cells, we observed a shift from immunosuppressive to more immune-active phenotypes (**Figure 5B-D**). Combined TIG+aCD40 reduced suppressive TREM2^+^ macrophages (9% of total myeloid cells) by 3.2-fold compared with aCD40 (29% of total myeloid cells) and 2.5-fold compared with TIG alone (23% of total myeloid cells). In addition, aCD40 and TIG+aCD40 induced significantly higher abundance of macrophages, 4-fold and 3.4-fold, respectively, expressing antigen-presenting gene signatures (APC Macs) compared with TIG treatment (with only 5% of all myeloid cells). A similar pattern was observed for mature, activated CCR7^+^ dendritic cells (DCs), with substantially more in the aCD40 and TIG+aCD40 groups than in the TIG group. In contrast, the TIG group expressed substantially more CLEC9a^+^ cDC1 than either the aCD40 or TIG+aCD40 groups. Another change in myeloid cells was a prominent increase in mast cells in the combined treatment compared with aCD40 or TIG alone. As the efficacy of combined treatment was reduced upon NK depletion, we further clustered NK cells into six subsets defined by distinct transcriptional profiles (**Figure 5E-G; Supplementary Figure 5**). Treatment altered the relative representation of several NK-cell states, including subsets 1, 3, and 5. Subset 1, characterized by a transcriptional program associated with inflammatory signaling, cellular stress responses, and immune activation, represented 24% and 18% of NK cells following aCD40 and TIG treatment, respectively, and increased to 28% with combined TIG+aCD40 treatment. Subset 3, expressing CXCR6, characterized by genes associated with immune regulation, trafficking, and cell-cell signaling, was most abundant following TIG treatment (32%), compared with aCD40 (7%) and TIG+aCD40 (18%). Subset 5 exhibited a proliferative, cell-cycle-associated transcriptional program and represented 7% of NK cells after aCD40 treatment, decreased to 1% with T/I, and remained at 7% with TIG+aCD40 treatment. Together, these findings indicate that TIG and aCD40 differentially change the composition and transcriptional states of tumor-infiltrating NK cells.

**Figure 5.**
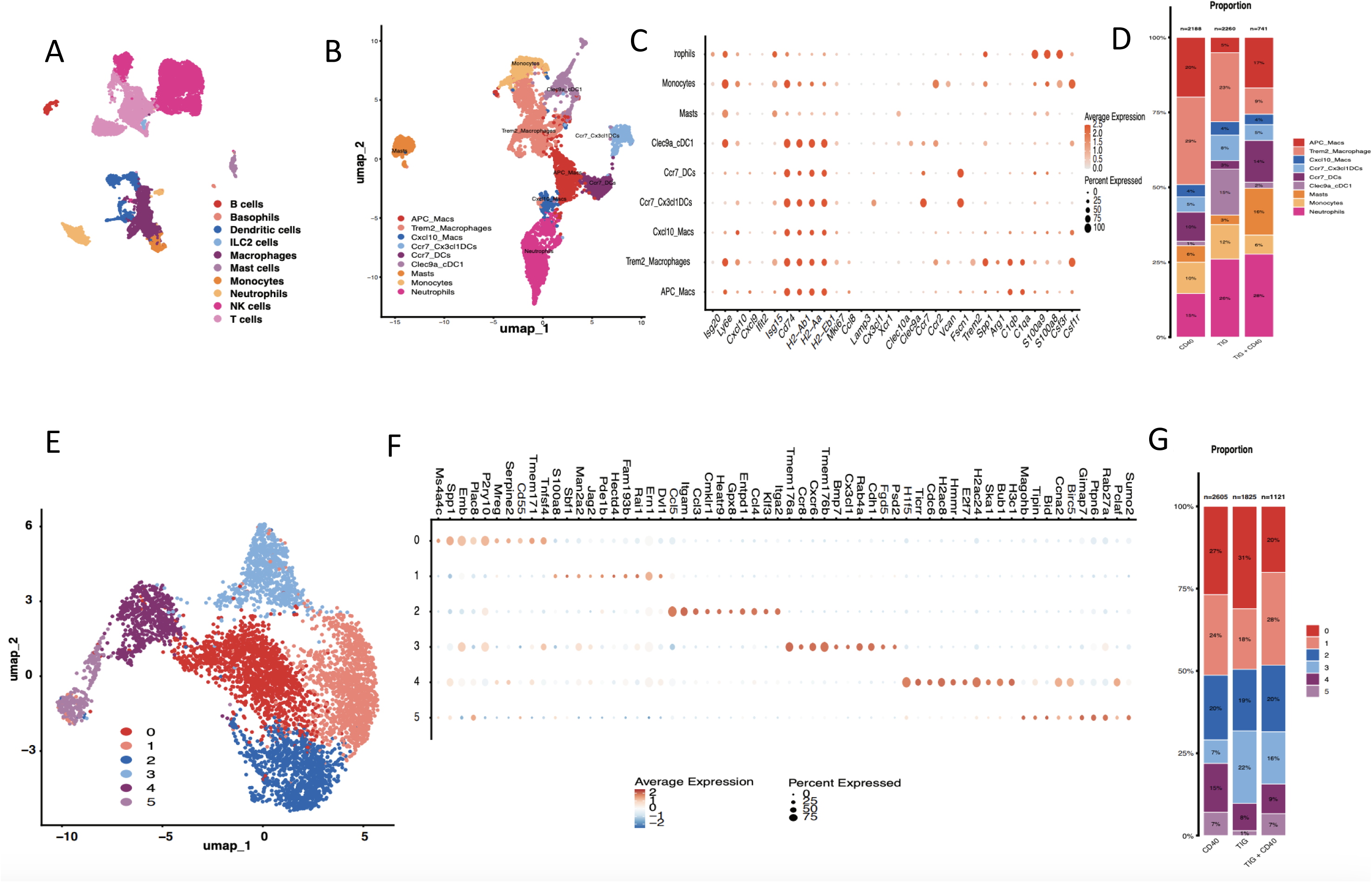
scRNA-seq analyses of immune cells in the 9464D-GD2 TME. Six days after the treatment started for NB-bearing mice, tumors were removed and analyzed by RNAseq. **A**. UMAP of immune cells, including all analyzed cells from the three treatment regimens, colored by identified cell type. **B**. UMAP of all myeloid cells analyzed from the three treatment regimens, colored by identified myeloid cell subtypes. **C**. DotPlot depicting the expression of marker genes used to define myeloid cell types. The color density of each dot indicates the average expression level of the marker gene; the dot size represents the percentage of cells that express the gene. **D**. Stacked bars illustrate the relative proportions of each myeloid cell subtype in each treatment regimen. **E**. UMAP of all NK cells analyzed from the three treatment regimens, colored by cluster identity for 6 separate NK subtypes. **F**. DotPlot depicting genes differentially expressed between the 6 identified NK cell subtypes. The color density of each dot indicates the z-scored expression level of the differentially expressed gene in each cell subtype, with the dot size representing the percentage of cells that express the gene. **G**. Stacked bars illustrate the relative proportions of each NK subtype in each treatment regimen.

### Absence of direct effect of aCD40 on 9464D-GD2 cells

Because some human NB cells express CD40, especially after stimulation with interferon gamma (IFN-γ),^29^ we tested if 9464D-GD2 cells expressed CD40, thereby potentially enabling aCD40 to have a direct effect on these NB cells. Flow cytometry shows that 9464D-GD2 cells do not express CD40 on their surface, even after stimulation with IFN-γ (**Supplementary Figure 6A**). **Supplementary Figure 6B** shows staining of CD40-expressing A20 B-cell lymphoma (positive control for aCD40 staining), and **Supplementary Figure 6C** shows upregulation of MHC-I expression on B16 melanoma cells stimulated with IFN-γ (positive control for IFN-γ function). These results demonstrate the absence of expression of CD40 on 9464D-GD2 and suggest that aCD40 does not kill 9464D-GD2 cells directly via apoptosis, a mechanism shown for some human NB cells.^29^ Furthermore, they are consistent with our data here showing the importance of immune cells (specifically macrophages and NK cells, **Figure 3**) in TIG+aCD40-induced antitumor responses for 9464D-GD2.

### Effect of TIG+aCD40 against multiple tumors in a minimal disease setting

Most patients with relapsed or refractory NB have multifocal disease, which can include measurable or very small multiple disseminated sites of disease.^30^ To simulate this multifocal disease setting, C57BL/6 mice were injected simultaneously with 9464D-GD2 cells i.d. into 3 locations: the left flank, the right flank, and the abdomen. Mice were randomized and treatment began when most of the tumors achieved ∼5 mm^3^ in volume. Whereas TIG alone did not result in any tumor regression (30 of 30 implanted tumors grew), consistent with what we showed in **Figure 1B**, TIG+aCD40 induced complete tumor regression (elimination of all 30 tumors) in all locations in all mice (**Figure 6A-D**). aCD40 alone was also effective, although to a lesser degree than TIG+aCD40. To test whether the tumor must be established to see the effect of TIG+aCD40, in a separate similar experiment, the treatment was started the day after implantation of 9464D-GD2 tumor cells in these same three locations in naïve mice divided into 2 treatment groups (TIG+aCD40 (n=5) vs. TIG alone (n=4)). As we showed with small established (∼ 5mm^3^) tumors (**Fig 6 A-D**), a combination of TIG and aCD40 rendered all 5 mice tumor-free in all three locations, while all 4 of the TIG-treated mice died of progressive disease, with 4 of 4 tumors implanted in the abdomen growing progressively, and 7 of 8 tumors implanted in either flank growing progressively (data not shown).

**Figure 6.**
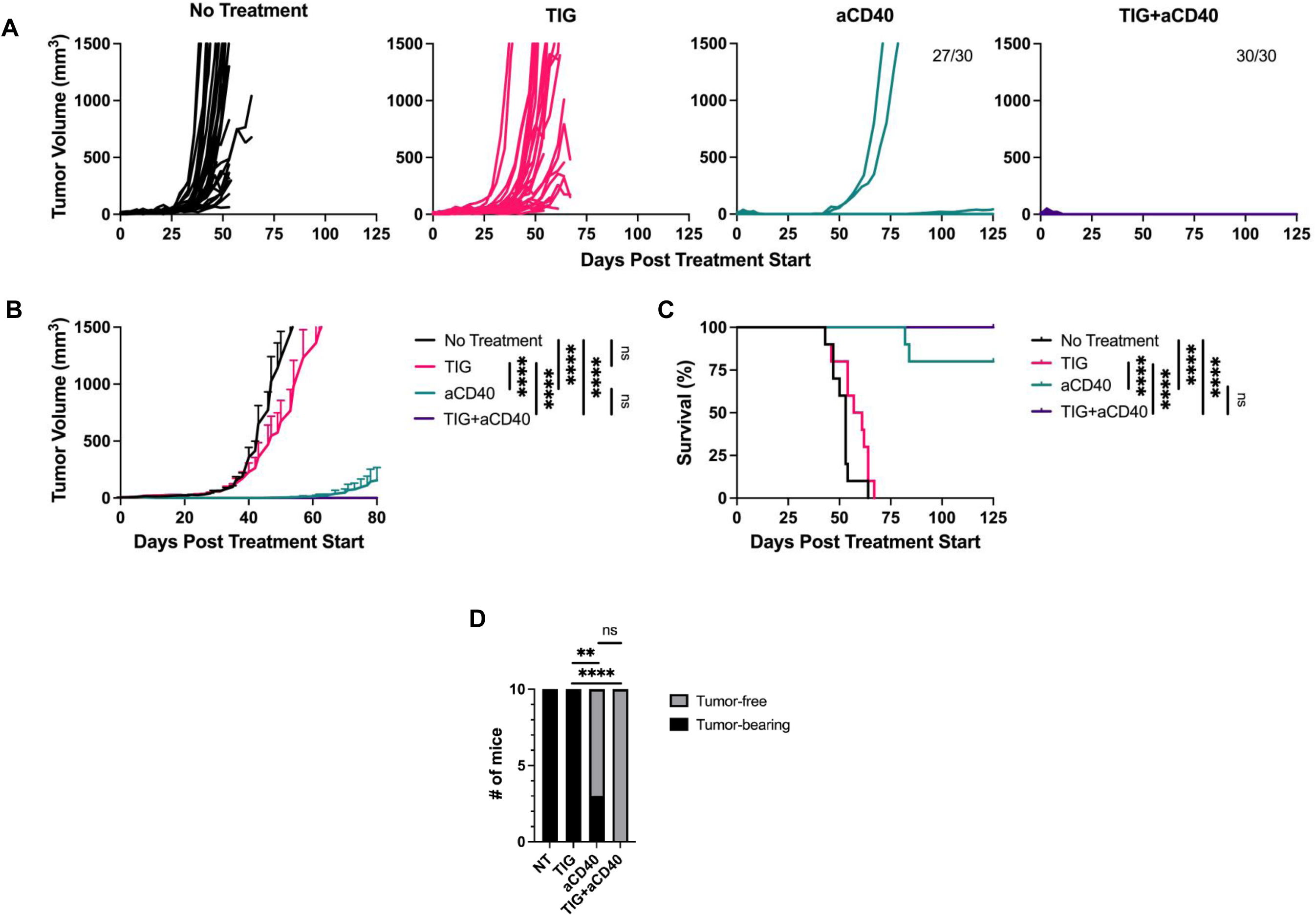
Effect of TIG+aCD40 against multiple small 9464D-GD2 tumors. C57BL/6 mice were injected i.d. with 9464D-GD2 tumor cells in the right flank, left flank and abdomen. Mice were treated with TIG, aCD40, TIG+aCD40, or NT when the tumor average volume in two or three sites was ∼5 mm^3^. The results show the individual tumor volumes (**A**), Mean tumor volumes +/-SEM (**B**), survival (**C**), and number of tumor-free mice at the end of two combined experiments (**D**), with 5 mice per group in each experiment (n=10 mice per group in the two combined experiments). **A** and **B** show all 30 tumors per group at each tumor implantation site in two experiments combined. Numbers in **A** indicate the number of tumor sites that remain tumor-free (if any) out of the total of 30 tumor implantation sites per group. **C** and **D** show the results of 10 mice per group in two combined experiments. \*\*\*\**p* < 0.0001.

### TIG+aCD40 treatment results in induction of T cell-mediated immunological memory

Because prevention of HR-NB relapse after treatment is clinically important, we tested if TIG+aCD40 results in immune memory. C57BL/6 mice cured of their 9464D-GD2 tumors (as in **Figure 1E,I**) were rechallenged ∼3 months later with the same dose of 9464D-GD2 cells in a new skin site. One group was treated with rat IgG, and another group received anti-CD4 + anti-CD8 mAbs to deplete T cells. The results show that mice cured with TIG+aCD40 either rejected the rechallenge tumors or suppressed their growth (**Figure 7A,B**), compared to the rapid growth of all tumors implanted into naïve mice. These rechallenged, previously cured, mice demonstrated extended survival (**Figure 7C**), indicating that they developed a memory response. This memory response was abrogated in T cell-depleted mice (**Figure 7A-C**), demonstrating that the memory response was T cell-dependent. In a separate similar experiment, nine C57BL/6 mice cured of their initial 9464D-GD2 tumors using TIG+aCD40 treatment were rechallenged with 9464D-GD2 NB implanted on the right side of the abdomen and syngeneic Panc02 pancreatic tumor implanted on the left side of the abdomen. Four of these nine mice rejected the 9464D-GD2 tumors, but none rejected the Panc02 tumors (**Figure 7D**), indicating that this memory immunity was tumor-specific.

**Figure 7.**
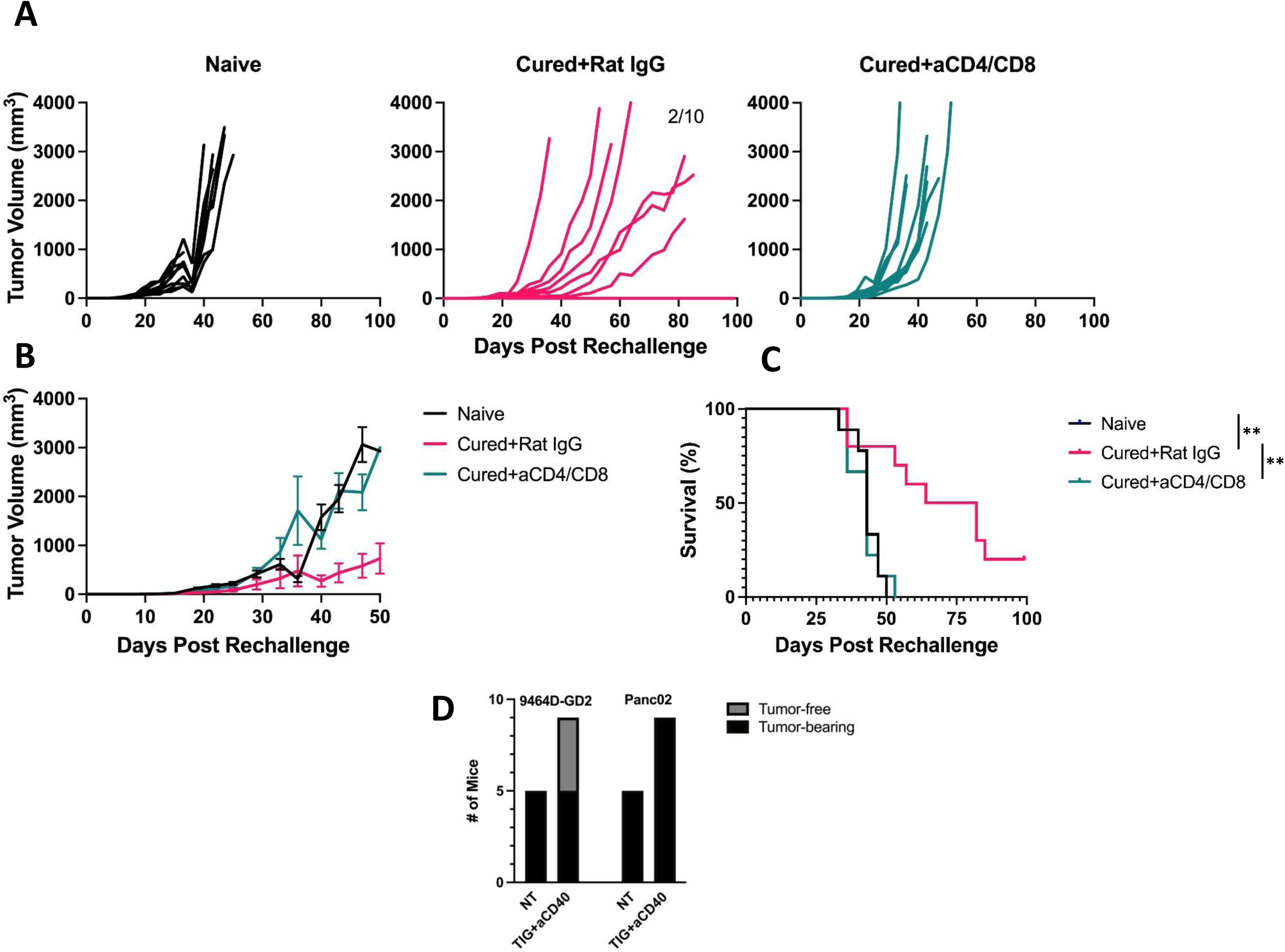
Immunological memory in mice that rejected their tumors following TIG+aCD40. **A-C.** After ∼3 months, mice that remained tumor-free (“cured”) after being treated with TIG+aCD40 were rechallenged with 9464D-GD2 cells in a new tissue site. Naïve mice were controls. Cured mice were injected i.p. with either a mixture of anti-CD4 and anti-CD8 mAbs to deplete T cells or rat IgG (control). The results show individual tumor volumes (**A**), mean tumor volumes +/- SEM (**B**), and survival (**C**) using all data from two combined experiments, 5 mice per group in each experiment (n=10 total mice per group). **D**. C57BL/6 mice bearing 9464D-GD2 tumors that became tumor-free following TIG+aCD40 treatment in two separate experiments (total n=9) were rechallenged on the right side of the abdomen with 9464D-GD2 cells and on the left side with Panc02 cells. Naïve mice (NT, n=5) were injected similarly as controls. The numbers of 9464D-GD2 and Panc02 tumor sites that developed tumors and those that remained tumor-free are shown.

## Discussion

TIG, a current clinical regimen for refractory/relapsed HR-NB, shows benefit as measured by a clinical response (partial or complete) in ∼40% of patients; however, this means ∼60% of children are resistant to this therapy, showing no response.^5,6^ Of the ∼40% of patients that do respond to TIG (full or partial response), without any further therapy only 25% remain progression-free after 4 years, and 75% go on to have disease progression.^31^ In other words, for HR-NB patients with relapsed/refractory disease, the current standard therapy provides 4-year progression-free survival for only ∼10% (40% of patients respond and 25% of these responders remain progression-free). The focus of the preclinical work reported here is on the development of a regimen that might build upon the TIG regimen to provide a better option for patients who do not respond to, or progress after, TIG, or potentially might be used in the future as an improved “first-line” regimen that is superior to TIG, for patients with relapsed or refractory disease. The preclinical work presented here provides data suggesting that the combined activation of NK cells and macrophages obtained by adding agonist aCD40 to TIG might have the potential to become one such improved regimen. We recognize that additional preclinical work (see below) is still needed before moving this concept into clinical testing. Even so, the findings reported here demonstrate that substantial therapeutic benefit can be obtained in mice bearing a tumor that is refractory to TIG when aCD40 is added.

The primary mouse tumor model in our studies, 9464D-GD2 NB, simulates most human HR-NBs as it is poorly immunogenic, MYCN driven, has low mutation burden, low expression of MHC-I,^21,23^ and is resistant to TIG. Our results demonstrate that adding agonistic aCD40 to TIG overcomes this *in vivo* resistance and induces complete tumor regression in most mice with smaller tumors. Importantly, cured mice show some resistance to tumor rechallenge with 9464D-GD2 tumors and demonstrate the development of T cell-mediated, tumor-specific, immunological memory.

For the treatment of 9464D-GD2 tumors in this setting, aCD40 used by itself was shown to have some antitumor activity (unlike TIG) but was not as effective as the combined TIG+aCD40 regimen. Our *in vivo* experiments, immune cell analysis in the TME by flow cytometry, and gene signatures by scRNA-seq demonstrate an important role for NK cells and macrophages in the antitumor effect of TIG and aCD40 against 9464D-GD2 NB. These results are consistent with our previous findings showing that the antitumor effect of aCD40 in several mouse tumor models was dependent on NK cells^16^ and macrophages.^17–20^ Importantly, scRNA-seq showed potentially distinct contributions of CD40 and TIG in inducing more effector NK cells and macrophages, indicating interplay between the two treatment components and two immune cell types that will be an important avenue for future studies. These results are also consistent with a study showing that IFN-γ-secreting mesenchymal stromal cells repolarized tumor-associated macrophages (TAM) and suppressed growth of human NB in nude mice; this antitumor effect was abrogated in macrophage-depleted mice.^32^ Importantly, TAM can also inhibit NK cells,^33^ the primary effector cells engaged by anti-GD2 mAbs to mediate ADCC.^34,35^ Our results show that both NK cells and macrophages play a role in the antitumor effects of TIG and aCD40 against 9464D-GD2 NB, highlighting the importance of interactions between macrophages and NK cells for this regimen of NB therapy.^36^

The precise cellular and molecular mechanisms of the antitumor effect of TIG+aCD40 are not yet clear. We’ve shown that TIG upregulated CD40 expression on peritoneal macrophages and that TIG+aCD40 decreased CD206^+^ M2 macrophages within TME. In addition, it is possible that TIG makes NB more accessible or susceptible to NK or macrophage-mediated killing^5–7^.

Although our results show a statistically significant reduction of the antitumor response in Csf1r-DTR/mCherry mice, it is important to note that macrophage depletion in those mice was not complete. Specifically, whereas macrophage depletion in the peritoneal cavity was ∼ 95%, it was ∼57% in the TME. Therefore, macrophages may play an even greater role in the TIG+aCD40-induced antitumor effect against 9464D-GD2 NB than we demonstrate in **Figure 3E-G**.

Our results show that neutrophil depletion resulted in statistically significant augmentation of the antitumor effect induced by TIG and aCD40. These data suggest that neutrophils in the TME of 9464D-GD2 tumors are predominantly immunosuppressive, similar to what we have observed with a separate immunotherapy approach.^19^ In various settings neutrophils can suppress T cells^37^ or NK cells and macrophages.^38,39^

Systemic aCD40 induces dose-limiting toxicity with several adverse side effects, which has been demonstrated in clinical trials.^13^ In our studies, systemically injected agonistic aCD40 did induce toxicity, as shown by transient weight loss and elevated liver enzymes in mice treated with aCD40 alone or TIG+aCD40. These findings agree with other preclinical studies^40,41^ reporting toxicity of aCD40. Therefore, there is a need to mitigate toxicity of aCD40 for use in NB patients. Reduction of aCD40-mediated toxicity may be achieved by several approaches, including: a) intratumoral injection of smaller doses as was shown by us^42^ and others;^43,44^ b) engineering the Fc portion of aCD40 to achieve enhanced binding to the inhibitory receptor FcγRIIB;^44,45^ c) use of aCD40 in bispecific antibodies;^46,47^ and d) prevention of aCD40-induced liver toxicity with an anti-TNF antibody administration.^48^ Some of these approaches already show clinical benefit.^45^

Our results show that 9464D-GD2 cells do not express CD40 even following stimulation with IFN-γ. Because some human NBs express CD40,^29^ TIG+aCD40 may exert some antitumor effect in the clinic for those CD40^+^ NBs not only by activated immune cells such as NK cells and macrophages, as shown here, but also via apoptosis induced by aCD40.^29^

The results presented here have several limitations. First, we have focused on a single tumor model. Extension of these findings to other mouse NB models (in other strains of syngeneic immunocompetent mice) and to human NBs growing in nude mice (which have intact NK cells and macrophages, the primary effector cells of TIG+aCD40 shown here for 9464D-GD2) is warranted. Second, we used an i.d. model of NB. It will be important to test the efficacy of TIG+aCD40 in a more clinically relevant orthotopic model where NB cells are injected into the adrenal glands. Third, patients with advanced, relapsed/refractory NB normally have observable or microscopic disseminated metastases. This TIG+aCD40 regimen should be tested in a murine model involving disseminated NB metastases. Using this model, we have established a premise for future work to investigate TIG+aCD40, including testing in separate murine models of HR-NB grown in different strains of mice, as well as both orthotopic models for NB disease and disseminated metastatic disease.^19^ Fourth, while we have shown that antitumor efficacy of TIG+aCD40 is more potent than that of either aCD40 alone or of TIG alone, we have not yet determined if each of the 3 components of the TIG regimen is required for the efficacy of TIG+aCD40, like we did when evaluating the role of each component of a separate, less clinically relevant aCD40-containing regimen.^49^ Fifth, as we have recently shown that aCD40 improves that efficacy of anti-GD2 mAb^19^, we will also consider whether adding T vs. I vs. TI further improves the efficacy of aCD40 + anti-GD2 mAb. Finally, for much larger 9464D-GD2 tumors, even though TIG+aCD40 can slow tumor growth, the tumors eventually all progress and the animals die. We need to further augment the efficacy of this TIG+aCD40 regimen. When aCD40+anti-GD2 mAb were combined with epigenetic modifier inhibitors, inclusion of checkpoint inhibitors (anti-CTLA4 and anti-PD1) improved the response to treatment.^19^ Additionally, a recent study by Menard *et al* in immunocompetent mice bearing a transplantable TH-MYCN NB showed that aCD40 mAb was far more effective when combined with anti-PD-L1 mAb (but not with anti-PD1 mAb), which depleted anti-inflammatory macrophages that preferentially express PD-L1.^50^ We are testing whether adding anti-PD-L1 can further augment the antitumor efficacy of TIG+aCD40 in mice bearing larger 9464D-GD2 tumors than those that can be cured by this TIG+aCD40 regimen. These above studies are beyond the scope of this current manuscript and most are underway, or planned, for inclusion in a follow-up report.

Overall, our results suggest that adding aCD40 might improve the efficacy of the current standard TIG regimen for relapsed HR-NB in children. Further preclinical testing is warranted to determine the best regimen and setting for potentially initiating clinical testing in relapsed/refractory NB in the future.

## Acknowledgements and grant support.

We thank Meridith Hyun for help with the statistical evaluation of the data and Gavin Fredericks for technical assistance. This work was supported by The Midwest Athletes Against Childhood Cancer (MACC) Fund, the University of Wisconsin Carbone Cancer Center and research grants from the Hyundai Hope on Wheels Foundation, the End Kids Cancer Foundation, The St. Baldrick’s Foundation, The Band of Parents Foundation, The Cure Childhood Cancer Foundation, The Children’s Neuroblastoma Cancer Foundation, Alex’s Lemonade Stand Foundation, The Super Jake Foundation, UWCCC Cancer Center Support Grant and by public health service grants R35-CA197078, and P01 CA250972 from the National Cancer Institute.

## Author contributions

AR: Conceptualization, Investigation, Writing –original draft, Writing – review & editing. JM: Data curation, Formal analysis, Investigation, Writing – review & editing. MF: Data curation, Investigation, Writing – review & editing. AH: Data curation, Formal analysis, Investigation, Writing – review & editing. AG: Data curation, Investigation, Formal analysis, Writing – review & editing. AU: Investigation, Formal analysis, Writing – review & editing. KD: Data curation, Investigation, Writing – review & editing. BH: Data curation, Investigation, Writing – review & editing. CC: Data curation, Investigation, Writing – review & editing. CW: Data curation, Investigation, Writing – review & editing. NH: Investigation, Data Curation, Writing – review & editing. JZ: Formal analysis, Software, Writing – review & editing. MS: Conceptualization, Methodology, Writing – review & editing. HD: Conceptualization, Funding acquisition, Investigation, Formal analysis, Writing – review & editing. AE: Investigation, Writing – review & editing. PS: Conceptualization, Funding acquisition, Project administration, Supervision, Writing – review & editing.

---

**Supplementary Figure 1.**
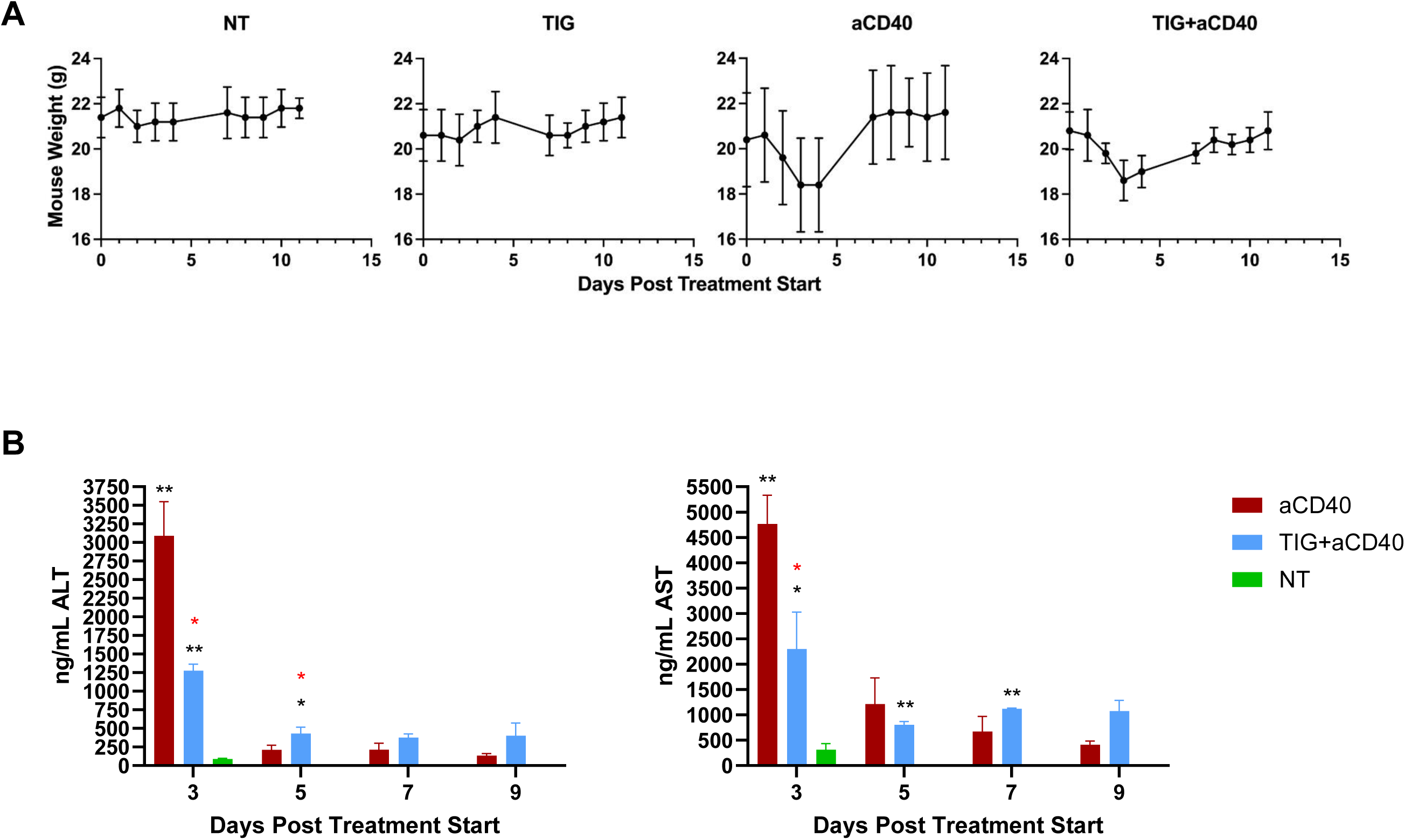
Effect of aCD40 and TIG+aCD40 on weight loss and liver enzyme levels. **A.** When aCD40, alone or with TIG, was injected into mice on day 1, they lost approximately 10% of their body weight over three days. This weight loss was transient, and mice regained their weight over the next 7 days. TIG alone did not induce weight loss. **B**. aCD40 and, to a lesser degree, TIG+aCD40 treatments induced a transient elevation of alanine aminotransferase (ALT) and aspartate aminotransferase (AST) on day 3 after the first TIG injection, 2 days after the aCD40 injection on day 1. Black asterisks show the differences compared to NT, and red asterisks show the differences between aCD40 and TIG+ACD40. No asterisks indicate no statistical significance. In separate experiments, TIG alone did not affect liver enzyme levels (not shown).

**Supplementary Figure 2.**
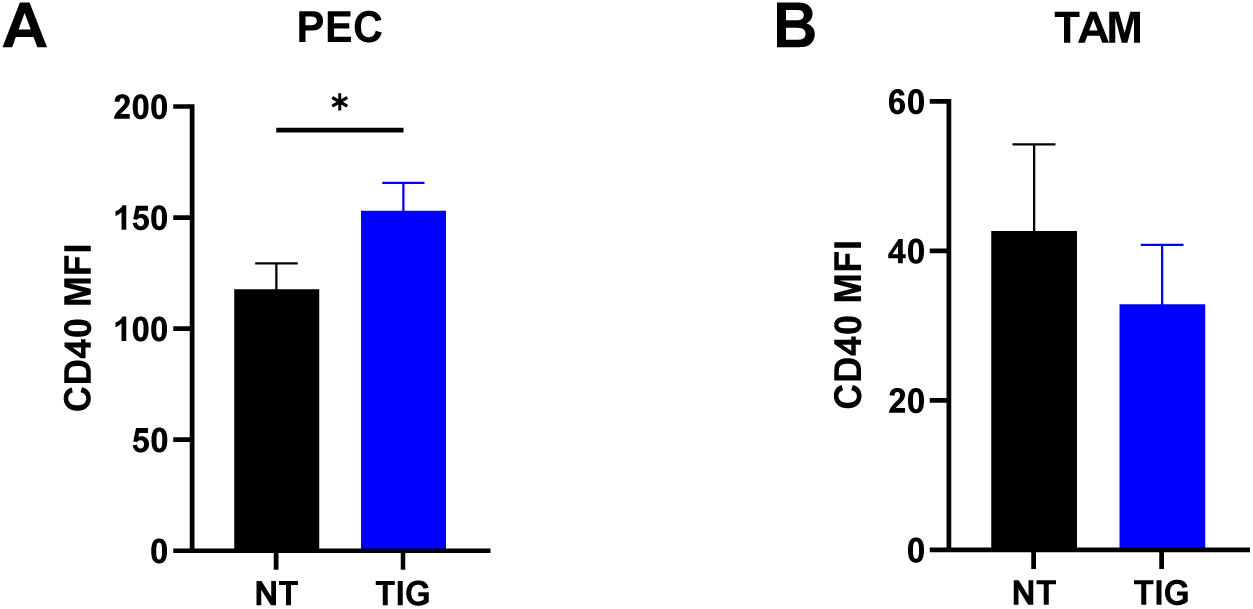
CD40 expression on macrophages after TIG injection. C57BL/6 mice bearing ∼ 50 mm^3^ 9464D-GD2 tumors were treated i.p. with TIG. PEC (**A**) and tumors (**B**) were collected on day 1 after the treatment. Means +/-SEM of CD40 expression on CD45^+^CD11b^+^F4/80^+^ cells are shown (n=3). Note: In the NT TAM group, the tumors were pooled from 3 mice to obtain enough cells for analysis. \**p* < 0.05.

**Supplementary Figure 3.**
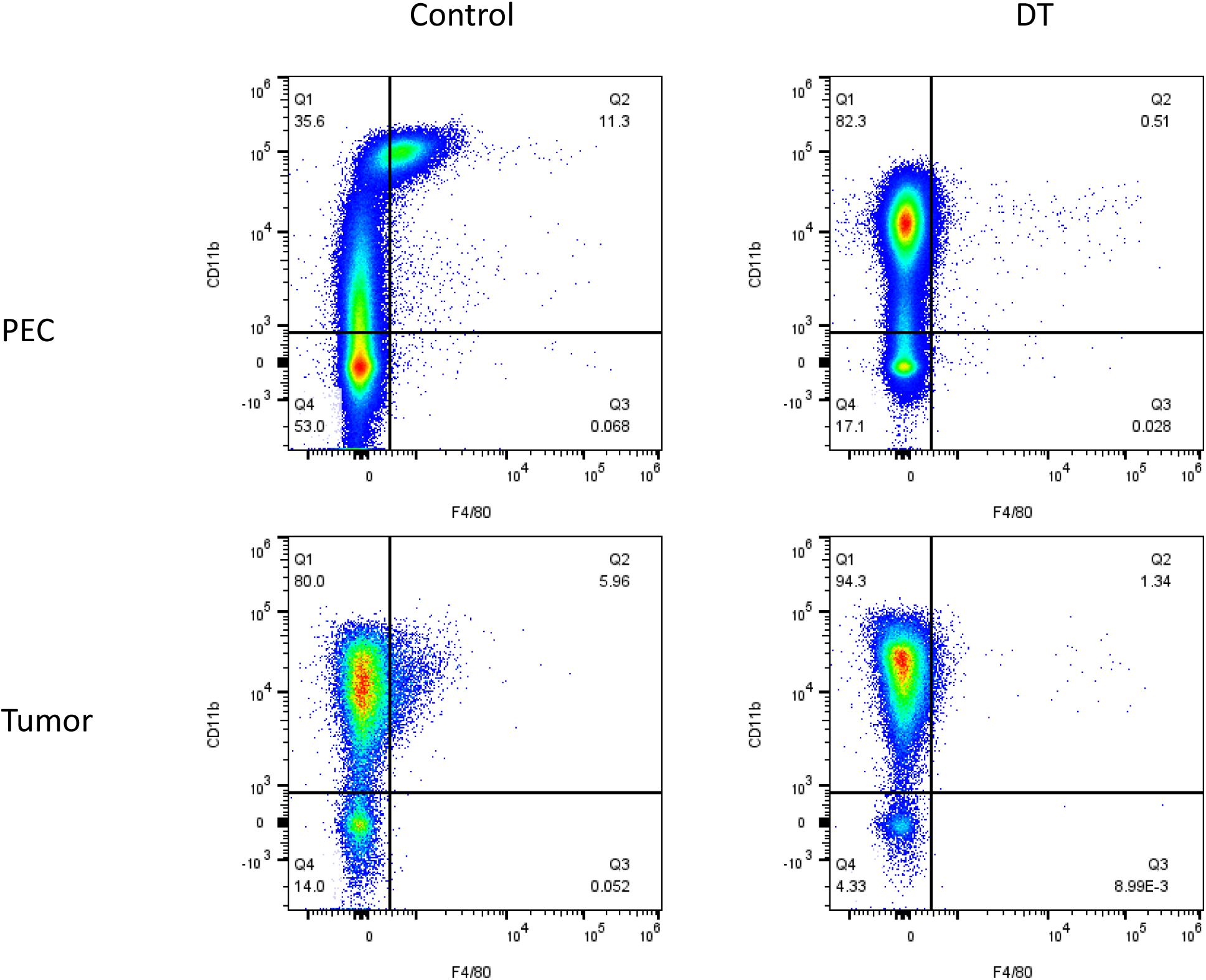
Efficacy of macrophage depletion in DTR-mCherry-*Csf1r* mice. DTR-mCherry-*Csf1r* mice bearing 9464D-GD2 tumors were injected with DT or PBS (control) on day -1, 1, and 3. Tissues were collected for flow cytometry on day 4. Representative dot plots of percentages of macrophages (CD11b^+^F4/80^+^) in PEC and tumors are shown for one control and one DT-treated mouse. When data from all 3 control and all 3 DT-treated mice were evaluated, macrophage depletion following DT administration was 95% in PEC and 57% in tumors.

**Supplementary Figure 4:**
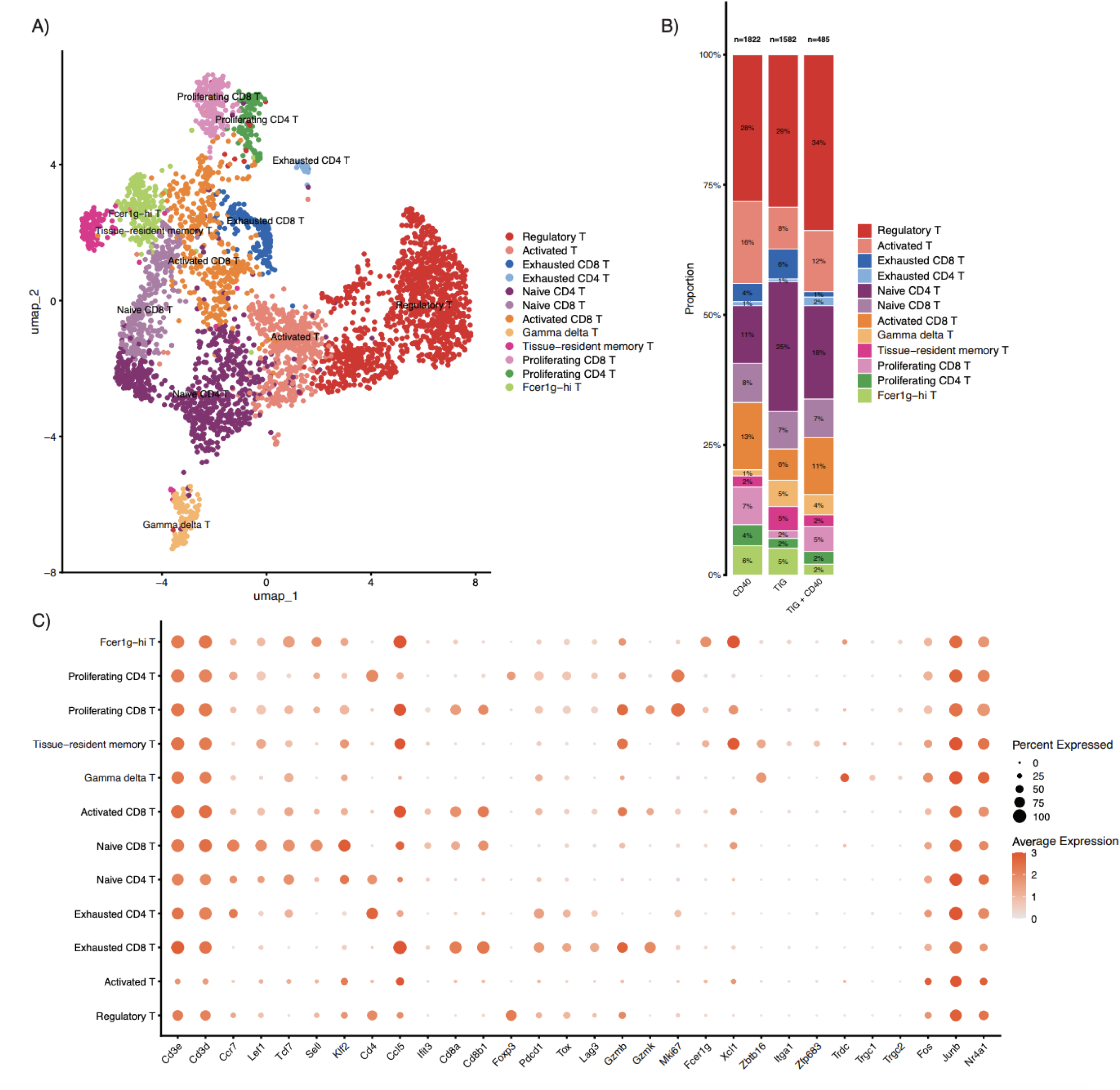
scRNA-seq of T cell subtypes. **A.** UMAP of all T cells analyzed from the three treatment regimens, colored by identified T cell subtypes. **B.** Stacked bars illustrating the relative proportions of the identified T cell subtypes in each treatment regimen. **C.** DotPlot depicting the expression of marker genes used to define T cell subtypes included in **A** and **B**. The color density of each dot indicates the average expression level of the marker gene in each T cell subtype, while the size of the dot represents the percentage of cells within the cell type that express the gene.

**Supplementary Figure 5:**
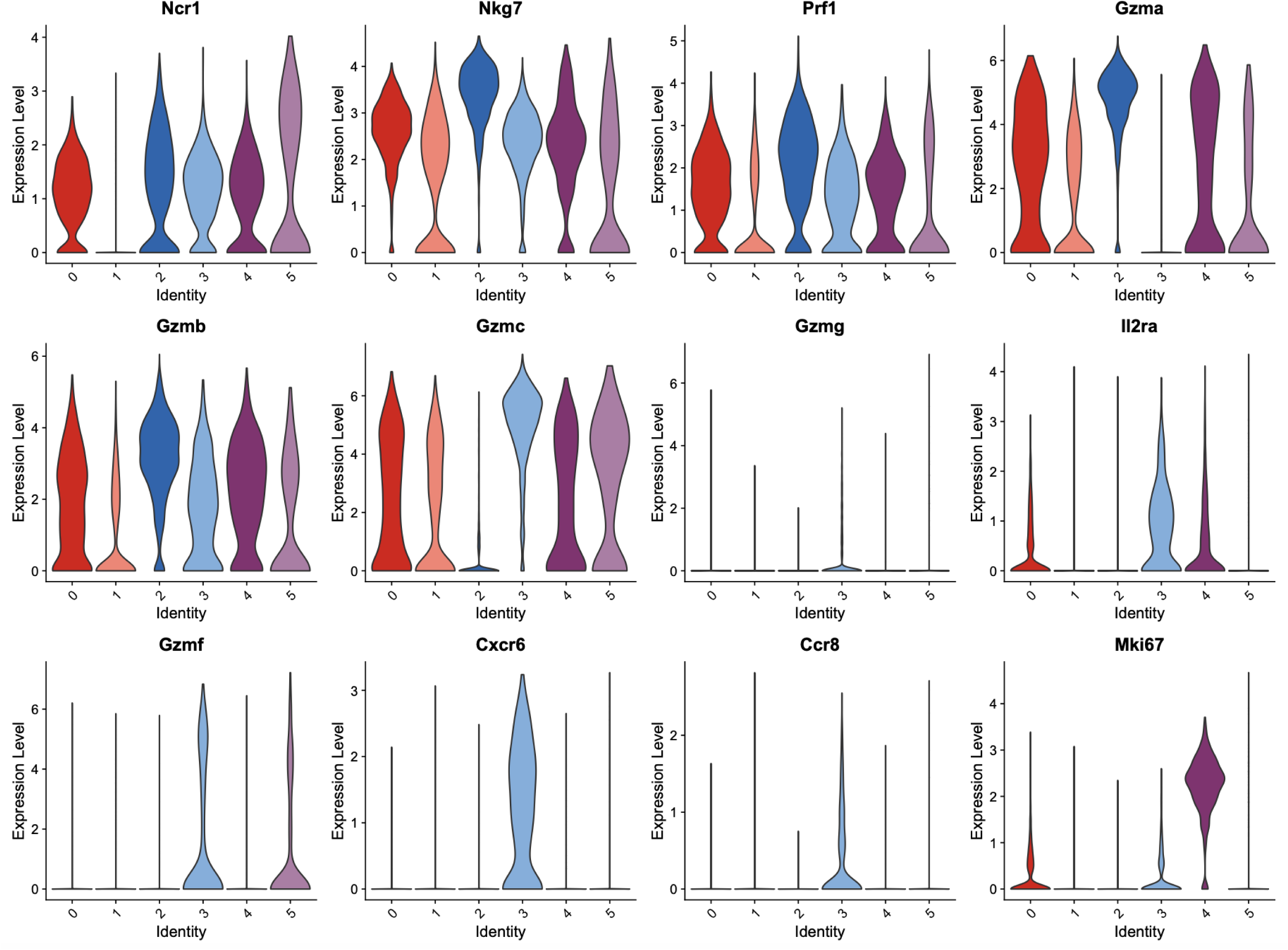
scRNA-seq of NK cell subtypes. Violin plots are shown representing the expression of key NK cell genes within the identified 6 separate NK cell subtypes.

**Supplementary Figure 6.**
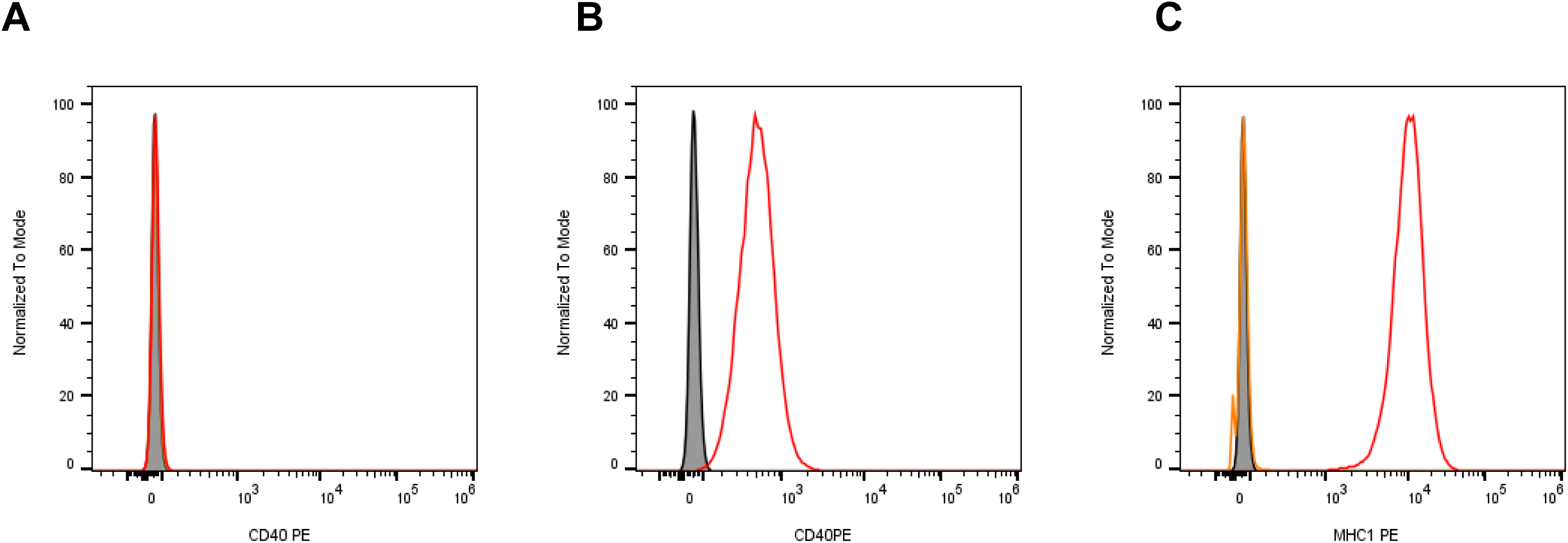
Absence of detectable CD40 expression on 9464D-GD2 cells. **A.** 9464D-GD2 cell line. Gray filled histogram=unstained cells, MFI=58.8; orange line=untreated cells, MFI=61; red line=IFNγ 100 U/mL for 48hrs, MFI=62.7. **B**. Positive control for CD40 expression, cell line A20. Gray filled histogram=unstained cells, MFI=57; red line=CD40 PE, MFI=513. **C**. Positive control for IFNγ stimulation, MHC class I expression in B16-F10 cell line. Gray filled histogram=unstained cells, MFI=59.9; orange line=untreated cells, MFI=61.6; red line=IFNγ 100 U/mL for 48hrs, MFI=10,123.

## Supplementary Materials and Methods

### Flow cytometry

C57BL/6 mice bearing 9464D-GD2 NB were treated with TIG +/-anti-CD40 as described above. On day 5 unless stated otherwise, tumors were harvested, cut into small pieces, and incubated for 30 minutes at 37°C in dissociation solution containing RPMI supplemented with 10% FBS, 1mg/ml Collagenase type D and 100µg/ml DNase I (Sigma-Aldrich, St. Louis, MO), using a Miltenyi gentleMACS Octo Dissociator. Samples were then incubated with Tonbo Ghost Dye Violet 510 for 30 min at 4°C, followed by washing in flow buffer (PBS-1% FBS) and a 10-minute incubation with anti-CD16/32 FC Shield (Tonbo, Clone 2.4G2) at 4°C. Cells were surface stained in Brilliant Stain Buffer (BD Biosciences) for 30 minutes at 4°C with CD45 FITC (30-F11), CD25 BB700 (PC61), CD8a APC-Fire 750 (53-6.7), CD4 BV785 (GK1.5), GD2 APC (14G2a), NK1.1 APC (PK136), GR-1 APC-R700 (RB6-8C5), MHC II APC-Fire 750 (M5/114.15.2), CD11b V450 (M1/70), Ly-6G BV711 (1A8), CD206 PE (MR6F3), F4/80 PE-Cy7 (BM8), Ly-6G FITC (S19018G), CD11b APC (M1/70), CD206 AF700 (C068C2), CD40 BV421 (3/23), CD19 BV711 (6D5), F4/80 PE (BM8), CD3 PE-Cy7 (17A2), F4/80 APC-Fire 750 (BM8), NK1.1 BV605 (PK136), CD40 PE (FGK45), and/or MHC I PE (34-2-12) according to the panel being used; these antibodies were obtained from BD Biosciences, BioLegend, or Invitrogen. Samples were washed, then fixed and permeabilized for 30 minutes at 4°C with the FoxP3/Transcription Factor Staining Buffer Set from eBioscience or the Cyto-Fast Fix/Perm Buffer Set from BioLegend following kit instructions. Samples were then stained with FoxP3 PE-Cy7 (FJK-16s) for 30 minutes at room temperature. All samples were then washed in permeabilization buffer and resuspended in flow buffer for acquisition on an Attune NxT flow cytometer (ThermoFisher). Data were analyzed with FlowJo v10.9.0 software (BD). Tregs were defined as live, single, CD45+ CD4+ CD25+ FoxP3+ cells and macrophages as live, single, CD45+, CD11b+, F4/80+ cells.

### scRNA-seq sample collection, library preparation, and sequencing

C57BL/6 mice bearing 9464D-GD2 tumors were treated with TIG, aCD40 or TIG+aCD40. Six days after the treatment started, seven to ten tumors for each treatment group were removed and pooled. Tumors were dissociated using the mouse Miltenyi Tumor Dissociation Kit in DMEM media containing FBS and penicillin/streptomycin. Live cells from the dissociated single cell suspensions were isolated using Miltenyi’s Dead Cell Removal Kit. Immune cells were then enriched with the STEMCELL EasySep mouse CD45 Positive Selection Kit. Non-immune cells were added back in at a 1:4 non-immune:immune cell ratio for each treatment group. Samples were fixed overnight for 21 hours in a fixation buffer containing 4% formaldehyde per 10x Genomics Fixation of Cells & Nuclei for GEM-X Flex Gene Expression protocol (CG000782). Following sample fixation, 10x Genomics’s GEM-X Flex Gene Expression Reagents Kits for Multiplexed Samples were used for probe hybridization, GEM generation, and library construction per the user guide (protocol CG000691 Rev B). Samples were sequenced on Illumina’s NovaSeq X Plus at Novogene. Raw sequencing reads were processed by CellRanger (v.9.0.0), which was also used to perform sample demultiplexing, barcode processing, and read alignment to the GRCm39 murine reference genome. Downstream analysis of single-cell RNA sequencing data was performed in Seurat (v5.5.0). Doublets were removed using scDblFinder (v1.20.2). Cells with fewer than 200 or greater than 10,000 detected features were removed, as were cells with percentage of mitochondrial reads exceeding 15%. Batch correction was performed using Harmony (v2.0.5). Broad and fine cell types were annotated using corresponding marker genes. Single-cell RNA-seq data have been deposited at GEO and are publicly available (GEO code: GSE310495).

